# Circadian rhythms in sexual behavior and their influence on reproductive outcomes in mice

**DOI:** 10.1101/2025.11.05.686629

**Authors:** Sydney Aten, Chiara Blake, Evan Keister, Adam Veith, Emma Fishbein, Oscar Ramirez-Plascencia, McKenzi Thompson, Shivani Howe, Sidney Pereira, Yaniv Sela, Victor M Navarro, Clifford B. Saper

## Abstract

Circadian clocks coordinate mammalian reproductive physiology, and circadian misalignment (resulting from shift work, jet lag, etc.) is known to impair fertility. Despite the well-established links between clock function and reproductive success, it remains unclear whether male and female mice maintained under standard *ad libitum* feeding conditions exhibit circadian rhythms in the propensity for sexual behavior, and to what degree such timing influences reproductive outcomes. Using standardized mating paradigms in C57BL/6J mice, we identified a circadian rhythm in sexual behavior in both sexes, with peak sexual activity most often occurring at circadian time (CT) 13–16 and a trough at CT4–7. To test the functional significance of these rhythms, we conducted 1-hour mating trials across four cohorts of C57BL/6J mice with pairs of mice having either aligned (e.g. male CT16 peak and female CT16 peak) or misaligned (e.g. male CT16 peak and female CT4 nadir) sexual behavior phases and monitored mating outcomes via ultrasound. While pregnancies were almost as frequent across all four cohorts, the numbers of live offspring were significantly more frequent when both the male and female mated at their peak phases than when both mice mated at their troughs. Notably, mating specifically during the male’s behavioral peak increased the likelihood of successful delivery of pups, suggesting that male circadian phase is a key determinant of miscarriage vs successful birth. These findings establish a circadian rhythm in the propensity for sexual behavior under standard feeding and housing conditions and indicate that mating time—particularly relative to the male’s circadian peak—can influence reproductive success. This insight provides a foundation for translational studies that explore whether intercourse timed to the circadian rhythms of the couple might help to increase fertility chances.

## Introduction

The body’s master circadian pacemaker, the suprachiasmatic nucleus (SCN), plays a central role in coordinating reproductive physiology and behavior in both humans and rodents (Miller and Takahashi 2013; Aten et al. 2024). In females, the SCN regulates the menstrual (and estrous) cycle and the preovulatory luteinizing hormone (LH) surge, thereby ensuring proper timing of ovulation. Similarly, copulation itself induces a pattern of circadian prolactin surges, which is likewise under control of the SCN (Freeman and Neill 1972; Poletini et al. 2010). Circadian control of reproductive physiology is not limited to females: healthy men (Arnal et al. 2016) and male rodents (Esquifino et al. 2004; Auer et al. 2020) exhibit daily rhythms in testosterone.

Further evidence linking the circadian clock to reproductive function comes from studies showing that disruption of clock timing impairs pregnancy success across species (Mills and Kuohung 2019). In female mice, genetic dysregulation of core circadian clock genes alters estrous cycle length and LH surge timing, in addition to leading to deficiencies in embryonic implantation, maintenance of pregnancy, and parturition (Miller et al. 2004; Miller and Takahashi 2013). Similarly, repeated phase shifting of the light-dark (LD) cycle dramatically reduces pregnancy success (Summa et al. 2012). In male mice, global loss of *Bmal1* and *Clock* leads to infertility/subfertility (Alvarez et al. 2008; Liang et al. 2013; Dolatshad et al. 2006; Ratajczak et al. 2009). In humans, chronic circadian disruption, often due to shift work or jet lag, has been associated with irregular menstrual cycles, altered reproductive hormones, pre-term birth, miscarriage, and infertility (Cone et al. 1998; Labyak et al. 2002; Baker and Driver 2007; Lawson et al. 2011; Nurminen 1998; Scherbarth and Steinlechner 2010; Mahoney 2010) and reduced fertility (Fernandez et al. 2020; Bisanti et al. 1996). Moreover, *Clock* gene variants are associated with altered semen quality in men with idiopathic infertility (Shen et al. 2015; Zhang et al. 2012), and shift-working men show increased rates of oligozoospermia, reduced sperm count, and elevated infertility risk compared with non-shift workers (Demirkol et al. 2021; El-Helaly et al. 2010; Sheiner et al. 2002; Liu et al. 2020).

These findings underscore the necessity of proper circadian function for successful conception. However, nearly all prior studies in mice have examined reproductive outcomes following broad disruptions of the circadian system—such as clock gene deletions or repeated light:dark (LD) shifts—without testing whether sexual behavior itself is under circadian control. Because such manipulations can produce wide-ranging pleiotropic and environmental effects beyond the circadian system itself, the specific contribution of intrinsic behavioral timing remains unclear. A foundational study by Richter (1970) reported that successful mating in rats depends on both the presence of estrus and intact circadian rhythmicity in males and females, suggesting that sexual behavior itself is temporally gated by the circadian clock. While restricted feeding has been shown to alter the daily pattern of male mouse mating behaviors (Kukino et al. 2022), no study, to our knowledge, has examined whether male *and* female mice, housed under constant conditions and provided food *ad libitum*, display endogenous rhythms in sexual behavior.

Here, we tested whether C57BL/6J male and female mice exhibit circadian rhythms in sexual behavior and whether the timing of mating relative to such a rhythm influences conception and eventual live births. These findings raise the possibility that *<u>when</u>* sexual intercourse occurs—relative to each partner’s circadian phase— may be an underappreciated factor influencing fertility outcomes.

## Results

### Male mice exhibit a peak in sexual behavior between CT13-CT16

To determine whether C57BL/6J male mice exhibit circadian rhythms in sexual behavior propensity, we dark-adapted sexually experienced male mice and performed 20-minute-long mating sessions across 8 different circadian timepoints. Each male mouse was tested at four of eight circadian timepoints each six hours apart, with mating sessions randomized and spaced one week apart (Fig. 1A). Across all timepoints, 69–79% of males exhibited mounting behavior at CT13 or CT16, compared to only 27% at CT7 (Fig. 1B; p = 0.0092, Fisher’s exact test, CT16 vs CT7). Interestingly, 10 of the 40 males did not display any sexual behavior across their four testing sessions and were excluded from behavioral analyses. Mounting frequency showed a clear circadian modulation, peaking at CT16 (Fig. 1C; F_(7,114)_ = 2.098, p = 0.0492, one-way ANOVA). Tukey’s post hoc tests confirmed a significant difference between CT7 and CT16 (adjusted p = 0.0425), with maximal mounting at CT13 (Fig. 1C, blue inset; F_(2.583,42.19)_ = 2.31, p = 0.0983, fixed-effect ANOVA). Percent time spent mounting also varied with circadian time, peaking at CT16 and reaching a minimum at CT7 (Fig. 1D; F_(7,111)_ = 2.898, p = 0.0081, one-way ANOVA). Post hoc tests revealed significant differences between CT7 vs CT16 (adjusted p = 0.0075) Together, these data indicate that in our fixed 20-minute mating assay, male sexual behavior peaks between CT13 and CT16.

**Figure 1.**
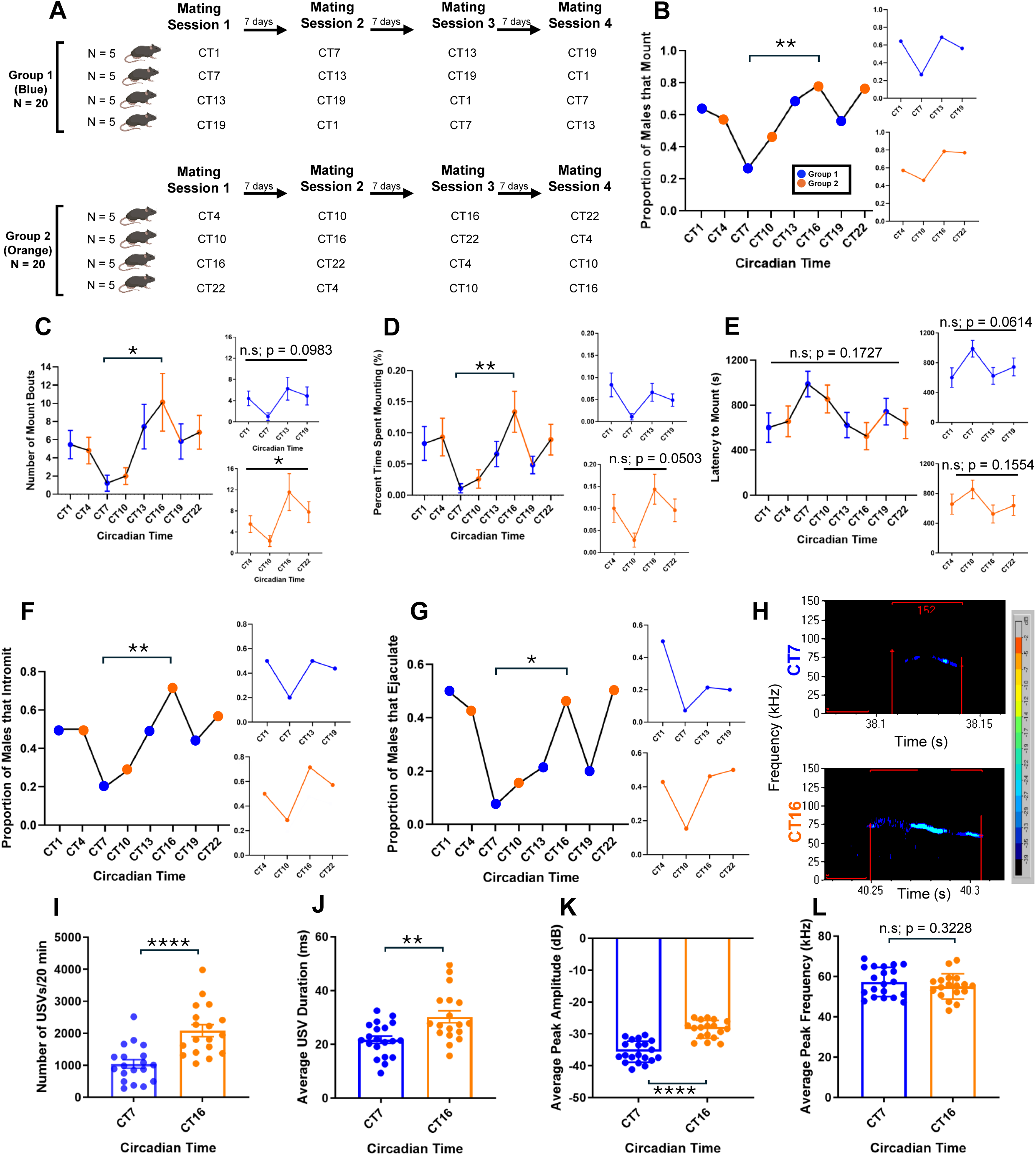
Rhythm in male mouse sexual behavior. A) Depiction of the experimental design where each male mouse was tested for mating behavior at CT1, 7, 13, and 19 (blue) with sessions spaced 7 days apart. A second cohort of males was tested at CT4, 10, 16, and 22 (orange). B) Proportion of all male mice that mounted within the 20-minute mating period. C) Number of mounts at each timepoint. D) Percent time spent mounting during each mating session. E) Latency to mount. F) Proportion of males to achieve intromission. G) Proportion of males to ejaculate. H) Representative sonograms of a single USV syllable emitted by males during a 20-minute session at CT7 and CT16. I) Average number of USVs. J) Average USV duration. K) Average USV peak amplitude. L) Average USV peak frequency. p < 0.05: *; p < 0.01: **; p < 0.0001: ****; n.s.: not significant. N = 13-20 mice per group.

Two other male consummatory behaviors, intromission and ejaculation, showed similar temporal patterns. Nearly 71% of males achieved intromission at CT16, compared to only 20% at CT7 (Fig. 1F; p = 0.0092, Fisher’s exact test, CT16 vs CT7). Moreover, 46% of males ejaculated during the 20-minute test at CT16, while only 7% did so at CT7 (Fig. 1G; p = 0.0329, Fisher’s exact test). A subset of males also ejaculated during the late circadian night and early circadian day (CT22–CT4; Fig. 1G). To assess whether this behavioral rhythm could reflect fluctuations in circulating testosterone, we collected serum from sexually experienced males across the same eight circadian timepoints (Supplementary Fig. 1A). Testosterone levels showed a modest peak at CT10, though this difference did not reach statistical significance (Supplementary Fig. 1B; F_(3.399,28.65)_ = 0.9381, p = 0.4444, fixed-effect ANOVA).

### Male mice display an increase in ultrasonic vocalizations (USVs) at CT16

Because male mice show ultrasonic vocalizations (USVs exceeding 30 kHz frequency) during sexual behavior (Nyby 1983; White et al. 1998; Holy and Guo 2005; Karigo et al. 2021) or in the presence of females or their urinary pheromones (Gourbal et al. 2004; Sipos et al. 1992), we next asked whether USVs follow a similar circadian pattern. As expected (White et al. 1998), during the mating sessions, our males produced numerous 40–70 kHz vocalizations (see Fig. 1H for representative spectrograms). Quantification revealed that the total number of USVs emitted per 20-minute mating session was significantly higher at CT16 than at CT7 (Fig. 1I; t_(35)_ = 4.597, p < 0.0001, unpaired t-test). Similarly, both the average USV duration (Fig. 1J) and peak amplitude (Fig. 1K) were significantly greater at CT16 (t_(36)_ = 3.350, p = 0.0019 and t_(36)_ = 7.011, p < 0.0001, respectively), while the average peak kHz frequency did not differ between the two timepoints (Fig. 1L; t_(36)_ = 1.003, p = 0.3228).Together, these results indicate that male mouse courtship vocalizations are enhanced at CT16, reflecting increased arousal and mating motivation during the active phase of the circadian cycle.

### Naturally cycling, estrus female mice show peak in receptive behaviors and delayed rejection behaviors at CT16

Female mice are sexually receptive only during the estrus (and occasionally proestrus) stage of the estrous cycle, when they are fertile. However, it is not known whether females exhibit greater receptivity at specific circadian times within the estrus period. To address this, we paired estrus-stage females with sexually experienced males for a 40-minute period at one of eight circadian timepoints (Fig. 2A). Estrous state was determined via vaginal cytology and visual inspection. As previously described (Byers et al. 2012), estrus females displayed an open, non-swollen vaginal opening (Fig. 2B, top right) and vaginal smears were dominated by cornified epithelial cells, in contrast to metestrus or diestrus females, which exhibited leukocyte-rich cytology and closed vaginal openings (Fig. 2B, left and lower panels).

**Figure 2.**
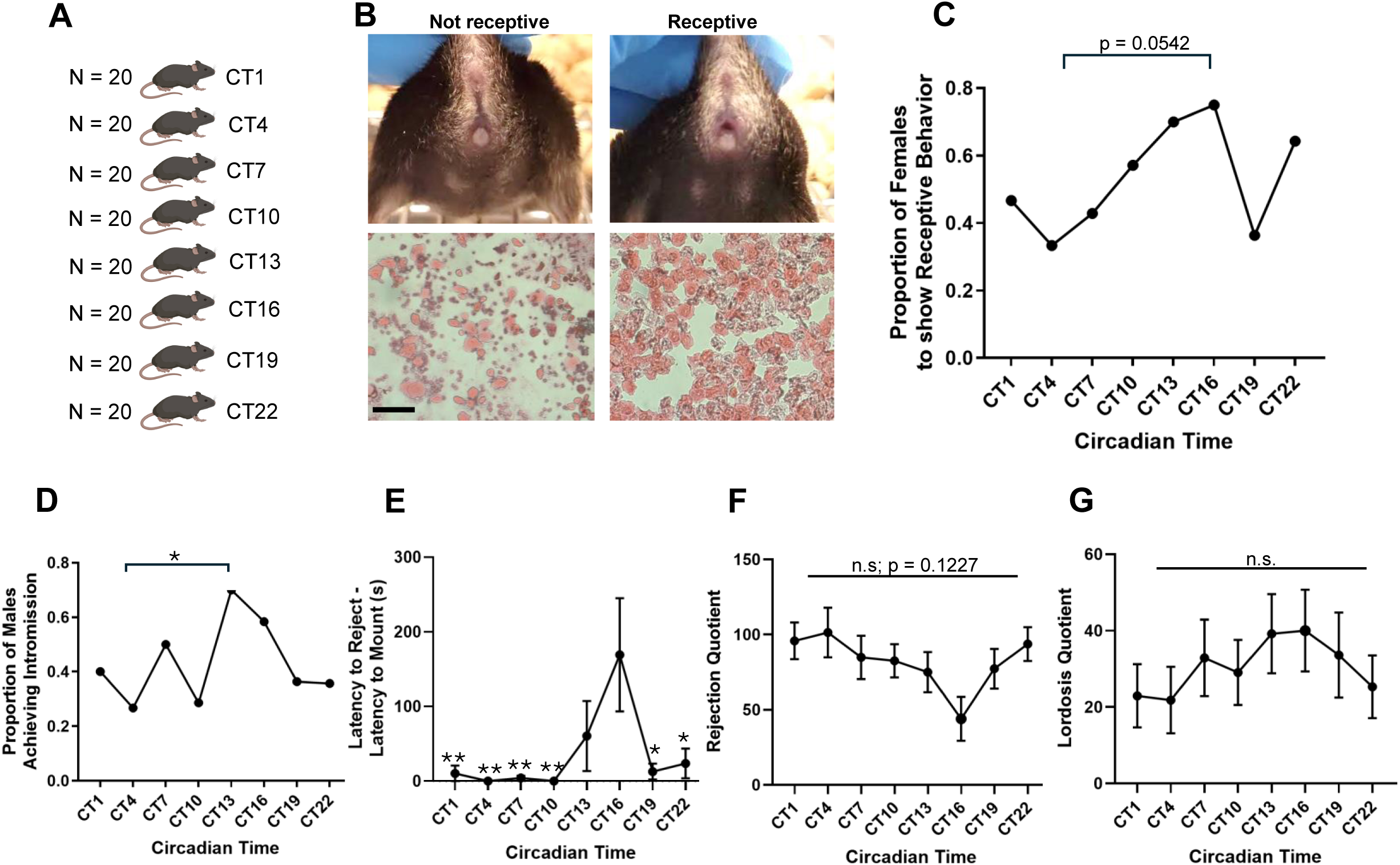
Rhythm in female mouse sexual behavior. A) Depiction of experimental mating paradigm in female mice. Female mice were only mated at one circadian timepoint. B) Representative images of a non-receptive (left, top panel) and receptive (right, top panel) female mouse. Note that the non-receptive female is in metestrus/diestrus given the closed vaginal opening and presence of leukocytes in the vaginal smear. The receptive female is in estrus given the open, non-swollen vagina and presence of cornified epithelial cells (and absence of leukocytes) in the vaginal smear. Scale bar = 100 µm. C) Proportion of females that showed receptive behavior, defined as having lordotic posture. D) Proportion of males that achieve intromission at the eight different timepoints. E) Latency for the female to reject. F) Rejection quotient, defined as rejection episodes/total mount attempts*100. G) Lordosis quotient, defined as lordotic episodes/total mount attempts*100. p < 0.05: *; p < 0.01: **; n.s.: not significant. N = 10-15 mice per group (after experiment completion).

We first examined the proportion of females displaying receptive behaviors. Approximately 75% of females were receptive at CT16, compared to only 33% at CT4 (Fig. 2C; p = 0.0542, Fisher’s exact test). Consistent with this, a greater proportion of females allowed males to achieve intromission at CT13 (70%) and CT16 (58%) than at CT4 (27%) (Fig. 2D; p = .0486, CT13 vs CT4 and p = 0.1302, CT16 vs CT4, Fisher’s exact tests), supporting enhanced sexual receptivity (Ventura-Aquino and Paredes 2023) between CT13 and CT16.

Next, we analyzed rejection behaviors (darting, kicking, or running from the male). A significant effect of circadian time was observed in the latency to reject (Fig. 2E; F_(7, 90)_ = 3.833, p = 0.0011, one-way ANOVA), with females at CT16 taking significantly longer to reject than those at CT1, CT4, CT7, CT10, CT19, and CT22 (Tukey’s post hoc tests, adjusted p < 0.05 for all comparisons). The rejection quotient (number of rejections divided by total mount attempts) did not differ significantly across timepoints, although a trend was observed for lower rejection at CT16 and higher rejection at CT4 (Fig. 2F; F_(7, 97)_ = 1.680, p = 0.1227, one-way ANOVA). The lordosis quotient (number of lordotic events divided by total mount attempts) also did not differ significantly across timepoints (Fig. 2G; F_(7, 97)_ = 0.5390, p = 0.8030, one-way ANOVA). These observations suggest that the attempts at resistance by the females at their nadir probably delayed mounting beyond the 40-minute test window in our assay in many cases, but the females who were willing to permit mounting during this time period did not further resist the male.

Taken together, these data indicate that female mice do exhibit increased receptivity and reduced rejection at CT16. This pattern temporally aligns with the male behavioral peak, suggesting that both sexes are most sexually active between CT13 and CT16, potentially facilitating synchronized mating and optimized reproductive outcomes.

### Sexual behavior peaks around CT14-15 in mice mated over a 6-day period

In our short-term (20–40 min) mating experiments, both male and female mice exhibited peak sexual behavior between CT13 and CT16. To determine whether this pattern persisted under more naturalistic conditions, we next examined mating behavior in pairs housed together for an extended period. Sexually experienced males and females were paired at CT1, CT6, or CT12 and recorded continuously for six days—long enough to capture at least one complete estrous cycle (Fig. 3A). All mounts, intromissions, and ejaculations were manually scored and timestamped (Fig. 3B) by a blind experimenter.

**Figure 3.**
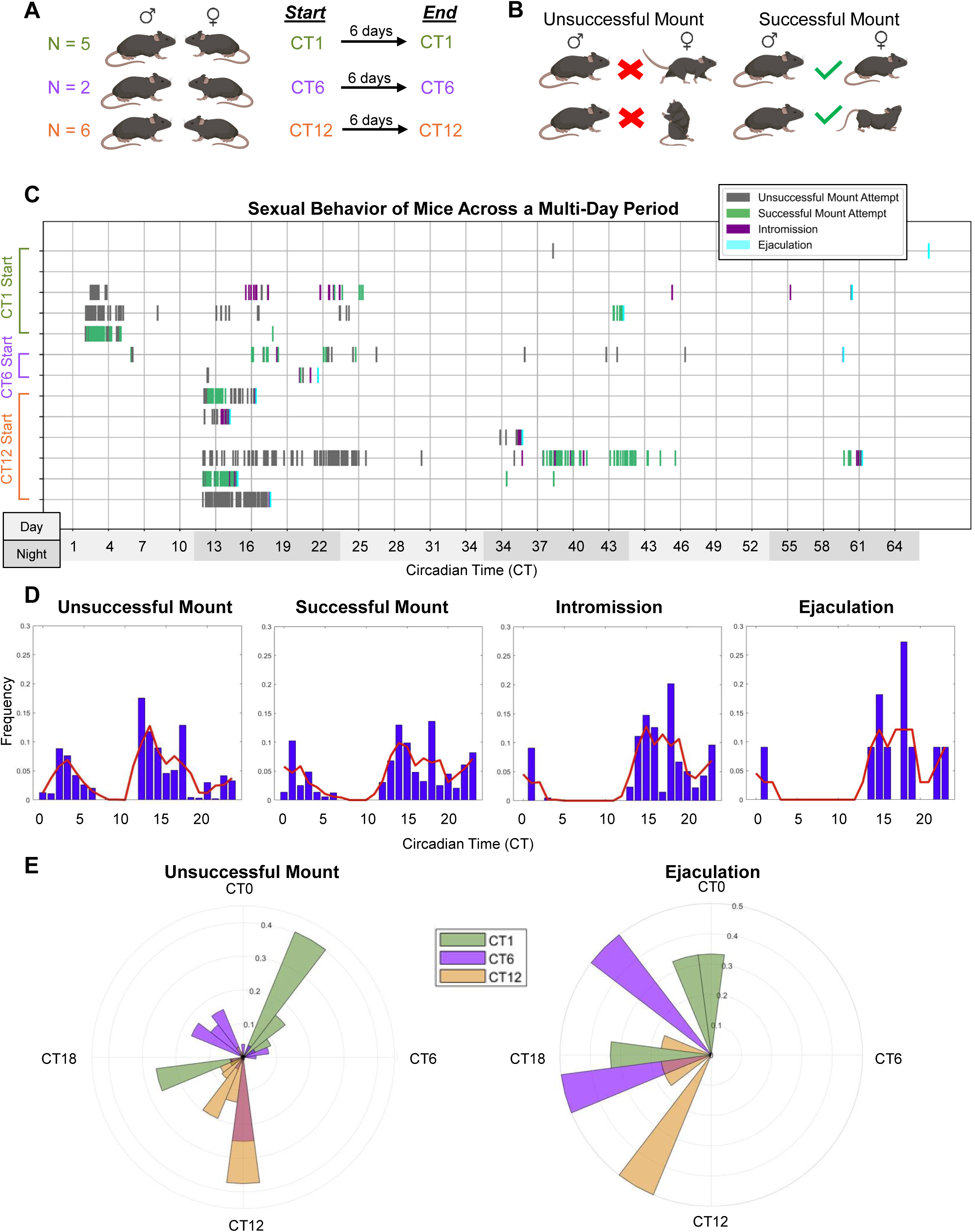
Long-term mating paradigm. A) 13 pairs of mice were mated, beginning at either CT1 (green), CT6 (purple), or CT12 (orange). Mice were in DD throughout the 6-day mating experiment. B) All mating events (e.g. unsuccessful mounts, successful mounts, intromission, and ejaculation) were timestamped. Unsuccessful mounts were defined as the female mouse running away, avoiding, or rearing/kicking the male. Successful mounts were defined by the female adopting lordotic posture or not running away from the male. C) Raster plot of mouse sexual behavior frequency recorded across the 6-day period. Note that the plot ends after 3 days given that mating activity had ceased in all groups after that time. The last recorded sexual behavior is indicative of male mouse ejaculation. Light gray bars on the X axis denote subjective day time periods, while darker gray bars denote subjective nighttime periods. D) Bar graph representation of all mice and their respective sexual behaviors. The red line shows the estimated probability of each event occurring at different circadian times. E) Rose plots of unsuccessful mounts (left) and ejaculation events (right). For the bar graph and rose plot graphs, for each animal, event times were binned into 24 hourly bins and normalized to yield a probability distribution (sum = 1). Normalized distributions were then averaged across animals to obtain the population-level temporal probability profile. N = 2-6 mating pairs per timepoint (e.g. CT1, CT6, CT12).

Across pairs, sexual activity occurred intermittently throughout the recording period, reflecting variation in female cycle stage. However, when behavioral events were plotted relative to circadian time, both intromission and ejaculation frequencies were found to cluster around CT14-CT15 (Fig. 3C-3D), consistent with the behavioral peaks observed in the short-term (e.g. 20 and 40-minute) assays described above (Fig. 1-2). The timing of pairing (e.g. CT1 vs CT6 vs CT12) had no significant effect on the time of day in which ejaculation occurred (p = 0.4366 Kruskal-Wallis test). No mating behavior (unsuccessful or successful, e.g. mounting resulting in copulation) occurred between CT7 and CT10 (Fig. 3D-3E).

### Circadian timing of mating impacts reproductive success

In our previous experiments (Figs. 1–3), we found that male and female C57BL/6J mice exhibit the highest propensity for sexual behavior between CT13–CT16 and the lowest during the circadian day. To determine whether the timing of mating affects reproductive success, we designed a one-hour long experiment to align or misalign the circadian phases of mating pairs (Fig. 4A). In group 1, both males and females were mated at their behavioral peak (CT16). In group 2, males were mated at their peak and females at their nadir (CT4). In group 3, males were mated at their nadir (CT7) and females at their peak (CT16), and in group 4, both sexes were mated at their nadir. If successful copulation occurred, pregnancy was assessed by ultrasound 6–7 days post coitum.

**Figure 4.**
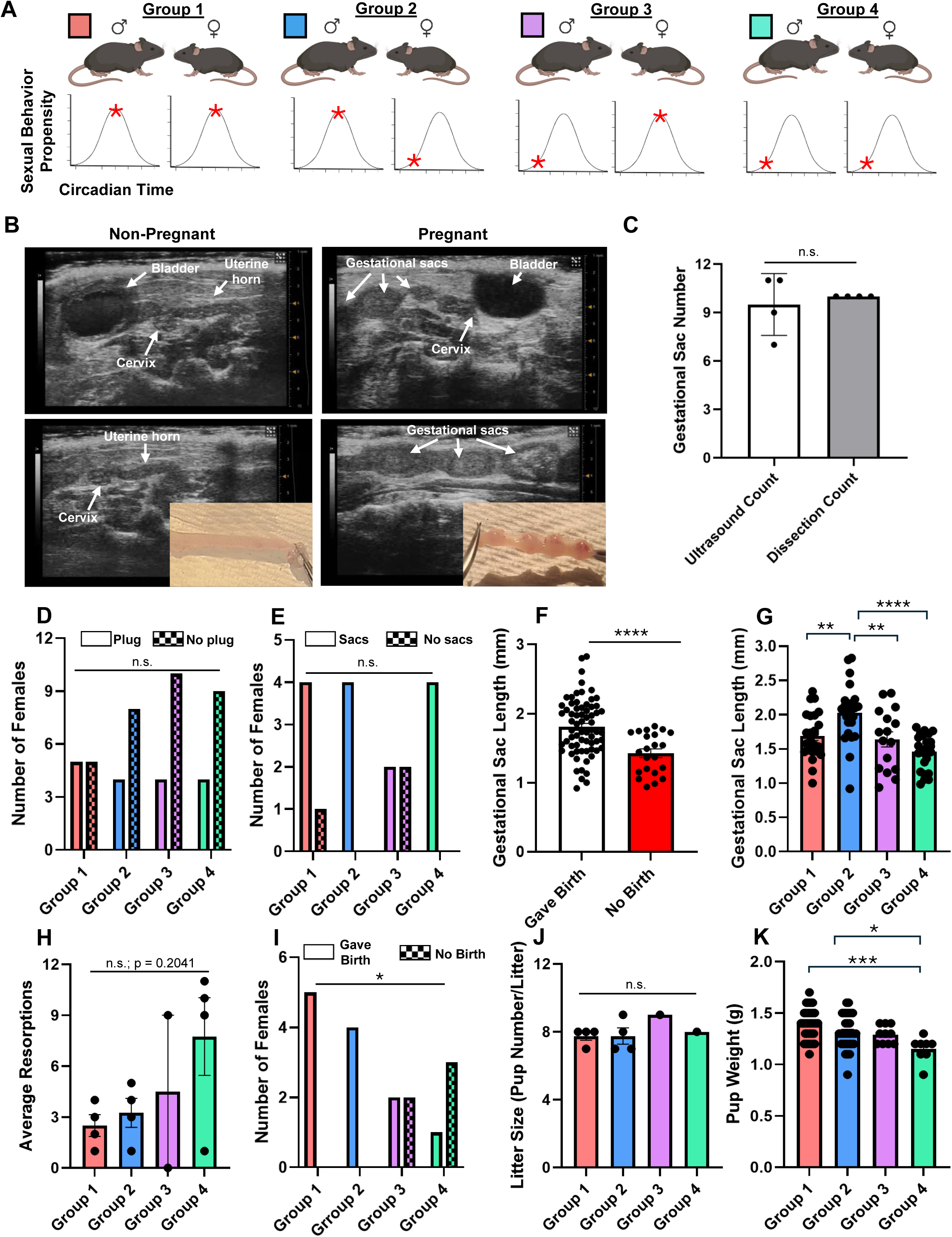
Circadian timing of mating impacts reproductive success. A) Schematic of mating alignment and misalignment paradigm. Mice were divided into four groups. In group 1 (red), males at their sexual behavior peak (e.g. CT13-CT16) were mated with females at their sexual behavior peak (e.g. CT16). In group 2 (blue), males at their sexual behavior peak were mated with females at their sexual behavior nadir (e.g. CT4). In group 3 (purple), males at their sexual behavior nadir (e.g. CT7) were mated with females at their sexual behavior peak. In group 4 (green), both males and females were mated at their sexual behavior nadirs. B) Representative images of an ultrasound from a non-pregnant mouse (left panels) and a pregnant mouse (right panels). The pregnant mouse was imaged at 6 days post-coitus. Note the gestational sacs within the uterine horns. Immediately after ultrasound imaging, mice were sacrificed and uterine horn contents were examined. As expected, no embryos were observed in the non-pregnant mouse upon dissection, while embryos were located in the pregnant mouse. Images depict a single uterine horn from each mouse. The brightness and contrast of the images was adjusted to better depict the horn and its contents and to visualize the difference between the two. C) Gestational sac number determined by ultrasound and subsequently by uterine horn dissection. D) Number of females in each of the four groups to have a copulatory plug after the 1-hour long mating experiment. E) Number of females in each group with observed gestational sacs at the 6–7-day post coitum ultrasound. F) Gestational sac length in females that failed to give birth compared to females that gave birth to pups. G) Gestational sac length at 6-7-days post coitum. H) Average resorptions (difference between initial gestational sacs observed at ultrasound and pups born). I) Number of females in each group that either showed visible gestational sacs at the 6-7 day ultrasound and gave birth or that showed visible gestational sacs at the 6-7 day ultrasound but that did not have live births (no birth). J) Average litter size (pups/litter). K) Average pup weight at time of birth. p < 0.05: *; p < 0.01: **; p < 0.001: ***; p < 0.0001: ****; n.s.: not significant. N = 4-5 females with copulatory plugs per group.

To validate use of ultrasound to determine pregnancy status, we performed ultrasound imaging on 4 female mice at six days post coitum, to test whether gestational sacs could be identified in the uterine horns (Fig. 4B, right), compared to non-mated (and thus non-pregnant) female mice (Fig. 4B, left). Immediately after imaging, these female mice were euthanized, and the uterine horns were dissected to examine the contents (e.g. to check for embryos). The number of embryos from each female were counted to determine accuracy between ultrasound gestational sac estimation and actual embryo number. We found no significant difference between gestational sac count on ultrasound and actual embryo count at dissection (Fig. 4C; t_(3)_ = 0.5222, p = 0.6376; paired t-test), confirming the reliability of ultrasound-based assessments.

Before examining pregnancy outcomes, we tested whether mating time influenced initial copulatory success. Although a greater proportion of females from group 1 (male peak–female peak) had a copulatory plug after the 1-hour long mating session, this difference was not statistically significant across groups (Fig. 4D; p = 0.7475, Fisher’s exact test). Similarly, the proportion of females with visible gestational sacs at 6–7 days post coitum did not differ significantly (Fig. 4E; p = 0.3765, Fisher’s exact test). Together, these data indicate that the circadian timing of sex did not have a significant effect on whether a copulatory plug and subsequent gestational sacs were observed. Out of 17 total females across all four groups that successfully mated (had a copulatory plug), only three (one from group 1 and two from group 3) showed no clear signs of gestational sacs during the live ultrasound.

To begin, we wanted to determine whether early measures of embryo development were associated with eventual pregnancy outcome. At 6-7 days post coitum, gestational sacs were significantly smaller in females that later failed to carry pregnancies to term (e.g. give birth) compared with those that ultimately gave birth (Fig. 4F; t_(88)_ = 4.028, p = 0.0001, unpaired t-test). To test whether early embryonic size could predict pregnancy outcome, we performed a logistic regression relating average gestational sac size to term pregnancy success. Average sac size significantly predicted pregnancy outcome (AUC = 0.76, 95% CI = 0.66– 0.86, p = 0.0003), such that smaller gestational sacs were associated with a higher likelihood of pregnancy loss. This finding is consistent with prior reports showing that resorbing embryos exhibit growth retardation (Flores et al. 2014; Peavey et al. 2017).

We next examined whether the timing of mating influenced early embryo development. Gestational sac size varied significantly among the four groups (Fig. 4G; F_(3,86)_ = 10.68, p < 0.0001, one-way ANOVA), with post hoc tests revealing significant differences between groups 1 and 2, 2 and 3, and 2 and 4 (p = 0.0071, 0.0059, and <0.0001, respectively). Females in groups 1 and 2 were significantly more likely to carry pregnancies to term (e.g. deliver pups) compared with those in groups 3 and 4 (Fig. 4I; p = 0.0394, Fisher’s exact test). Further, although the average number of resorptions (difference between initial observed gestational sacs and pups born) tended to be higher in groups 3 and 4, this difference did not reach significance (Fig. 4H; F_(3,10)_ = 1.838, p = 0.2041, one-way ANOVA). Among litters that were born, pup number did not differ significantly across groups (Fig. 4J; F_(3,6)_ = 0.8000, p = 0.5376, one-way ANOVA), but pup weights at birth were significantly affected by mating time (Fig. 4K; F_(3,77)_ = 6.212, p = 0.0008, one-way ANOVA), with pups from group 1 and group 2 weighing significantly more than those from group 4 (p = 0.0005 and p = 0.0316, Tukey’s post hoc tests).

Taken together, these findings suggest that the circadian timing of mating influences reproductive outcomes. Specifically, when both sexes mate during their circadian behavioral peaks, females are more likely to give birth (and not miscarry) and offspring are heavier at birth, indicating that alignment of male and female circadian rhythms enhances reproductive success.

## Discussion

Here, we show that male and female C57BL/6J mice display circadian rhythms in sexual behavior propensity, peaking between CT13 and CT16. Importantly, this behavioral rhythm has functional consequences as reproductive success was higher when males and females were mated during their respective behavioral peaks than when they were mated during their behavioral nadirs.

The timing of peak sexual behavior in males was not unexpected. The CT13–CT16 window coincides with the period of highest locomotor activity in mice (Todd et al. 2018; Abdollahi Nejat et al. 2024; Warfield et al. 2023; Kukino et al. 2022). Similarly, Kukino et al. (2022) in restricted feeding experiments observed increased mounting behavior in C57BL/6J males during the first six hours following food availability, when the mice are active.

An intriguing observation was that approximately 25% of males (10 out of 40) displayed no sexual behavior during any of the four mating sessions, in spite of the fact that they had been given two prior sexual experiences. While we did not determine whether these males might have preferred male partners, Karigo et al. (2021) reported that roughly 20% of males in their study exhibited male-directed mounting behaviors accompanied by USVs—patterns typically associated with female-directed mating. Our results, together with theirs, raise the possibility that this subset of males may exhibit homosexual or asexual behaviors. Future studies should examine whether these males display circadian rhythms in same-sex sexual behavior similar to those observed in typical heterosexual males.

In females, sexual receptivity also varied across the circadian cycle. Specifically, females took longer to reject male mounting attempts at CT16, suggesting a peak in receptivity at the same circadian phase as males. This ‘overlap’ between the sexes may reflect evolutionary pressure favoring temporal alignment in mating behavior to maximize fecundity. It also prompted us to ask whether reproductive success depends on the alignment between male and female circadian rhythms in sexual behavior. Along these lines, previous work has demonstrated that disrupting synchrony between the internal circadian clock and the external light–dark cycle can impair pregnancy outcomes. For example, Summa et al. (2012) found that repeated phase shifts reduced the proportion of pregnancies carried to term, particularly in female mice subjected to phase advances. Our current study is unique, however, as we show that negative reproductive outcomes can occur without external circadian disruption—simply by misaligning the internal sexual behavior phases of the mating partners. Indeed, we found that having males and females mate at their sexual behavior peaks (Fig. 1-2) improved pregnancy success and offspring outcomes, whereas misalignment reduced them (Fig. 4). To our knowledge, this is the first study to demonstrate that the circadian timing of mating itself can influence pregnancy success.

A particularly intriguing finding from our study was that pregnancy outcome depended strongly on the male’s circadian phase at the time of mating. Before we started, we had hypothesized that females mating at their peak would be more likely to sustain viable pregnancies, but that the circadian phase of the male might not be important for this outcome. Instead, females mated with males during their sexual behavioral peak were substantially more likely to carry pregnancies to term, regardless of the female’s sexual behavioral phase. In contrast, when males were mated at their behavioral nadir, pregnancy success dropped sharply—even when females were at their peak (Fig. 4). Notably, the number of gestational sacs detected by ultrasound at 6–7 days post coitum did not differ across groups, suggesting that fertilization and implantation may have occurred normally. Thus, the reduction in term pregnancies likely reflects increased post-implantation loss or early embryonic resorption in matings involving males at their circadian nadir.

These results point to the male’s circadian phase as a dominant factor shaping reproductive outcome, potentially reflecting circadian modulation of endocrine, gamete, or seminal factors. Although we did not directly assess these mechanisms, circadian variation in testosterone secretion and gonadotropin release has been documented in both rodents and humans (Ana I. Esquifino et al. 2004; Auer et al. 2020; Luboshitzky et al. 1997; Gall et al. 1979; Guignard et al. 1980), and androgens are well known to regulate spermatogenesis and sexual behavior (Smith and Walker 2014; Cunningham et al. 2012). Therefore, one could posit that males mating during their sexual behavioral peak may exhibit an optimal endocrine milieu that enhances sperm function or ejaculate composition. Supporting this idea, circadian *disruption* reduces sperm motility and acrosome reaction in rats (Travicic et al. 2023), and seminal plasma factors have been shown to modulate uterine receptivity and implantation (Bromfield 2014). Collectively, these findings suggest that males mating during their circadian peak may deliver seminal signals that are temporally optimized for fertilization and early embryonic development. However, the mating with males at their nadir did not result in fewer gestational sacs at day 6, although they were smaller. This suggests that the ability of the DNA in the sperm to produce a viable embryo may be affected by circadian phase. Of note, in other work in our lab, the body temperature of C57BL/6J male mice at CT4 is typically about 1.5° C cooler (around 36° C) than at CT16 (around 37.5° C). It is possible that sperm that are produced at the lower temperature may be less likely to produce a viable fetus, as is the case for ova used for *in vitro* fertilization which must be held closely at 37° C (Consensus Group 2020). Future studies that aim to identify this underlying mechanism are of great importance, as the timing of sexual behavior relative to the male’s internal clock may represent a critical yet underappreciated determinant of reproductive success.

Several limitations of our study should be acknowledged. First, we varied the duration of the window for measuring sexual behavior in our experiments to highlight the behavioral patterns. We used a 20-minute window to test male mice for their circadian sexual behavior, as this emphasized the sexual drive of the males (i.e., males with higher drive would perform earlier). For testing females, we used a 40-minute window, because they would not always have male behaviors to respond to in the first 20 minutes, and our preliminary experiments showed that they would often take longer to engage (sexually) with the male. While nearly all mice, if paired for a sufficiently long period, would likely produce pregnancies (or at least copulatory plugs), we wanted the testing of circadian phase of sexual behavior to emphasize the alacrity with which the partners would engage.

These mating methods identified rhythmic patterns in sexual behavior in male and female mice but did not determine their mechanistic basis. We observed a trend toward a circadian rhythm in serum testosterone in male mice, with a peak at CT10 (Supplementary Fig. 1)—several hours before the sexual behavior peak. The earlier peak in in testosterone compared to sexual behavior is consistent with the idea that some endocrine signals, particularly steroid hormones, often precede behavioral outputs and have delayed effects (Falkenstein et al. 2000). This anticipatory coordination of the endocrine system may prime male sexual motivation such that it occurs hours later in the active phase. Thus, even modest rhythmic fluctuation in testosterone may contribute to the timing of sexual behavior through downstream modulation of androgen-sensitive pathways in the hypothalamus. In addition, neural circuits within the hypothalamus may gate sexual behavior propensity such that male mice are most receptive between CT13-CT16. Ongoing studies in our laboratory are focused on delineating these neural circuit–based mechanisms.

Likewise, the mechanism underlying the female’s circadian rhythm, particularly in latency to reject, remains unclear. In ovariectomized, hormone-primed females, Kukino et al. (2022) reported that luteinizing hormone (LH) levels peak between ZT12 and ZT14, suggesting that rhythmic gonadotropin secretion could contribute to the observed behavior pattern. Although we did not measure LH in our naturally cycling females, it is plausible that a similar rhythm enhances receptivity at CT16. Indeed, circadian signals from the SCN are required to induce the preovulatory surge of gonadotropin-releasing hormone that triggers the LH surge (Miller and Takahashi 2013). Neural circuits within the hypothalamus likely also gate sexual receptivity in a time-dependent manner. To this end, Yin et al. (2022) identified cholecystokinin A receptor–expressing neurons in the lateral ventromedial hypothalamus that dynamically regulate female mouse sexual behavior. Whether such neurons exhibit circadian rhythmicity or receive direct or relayed SCN input remains an open question.

In summary, both male and female mice possess intrinsic circadian rhythms in sexual behavior parameters, with a shared peak during the early subjective night (CT13–CT16), coinciding with their active phase. The temporal alignment of these rhythms likely evolved to optimize reproductive success by synchronizing mating behavior with internal physiological readiness. Our findings further suggest that the male’s circadian phase plays a dominant role in determining pregnancy success, potentially via rhythmic modulation of endocrine or gamete factors, while female’s rhythms in sexual behavior likely ensures that sexual motivation and reproductive physiology coincide with the optimal window for fertilization. In this way, circadian control of sexual behavior in both sexes may serve as a finely tuned system that synchronizes mating with the most favorable internal conditions for conception, implantation, and pregnancy maintenance.

By characterizing a sexual behavior rhythm that predicts reproductive outcome, our work establishes a foundation for future studies examining how neural, endocrine, and gamete circadian oscillators interact to regulate fertility. Moreover, these findings may have potential translational implications: in humans, chronic circadian misalignment caused by night-shift work or irregular sleep schedules has been associated with infertility and pregnancy complications (Gamble et al. 2013). Our data raise the intriguing possibility that temporal alignment between partners’ biological clocks may influence conception success, highlighting the broader role of circadian biology as a critical—yet underappreciated—determinant of reproductive health.

## Materials and Methods

### Animals

All animal care and experimental procedures were approved by the Institutional Animal Care and Use Committee (IACUC) at Beth Israel Deaconess Medical Center. Mice were housed at 22 ± 1° C in a 12:12-hour light/dark (LD) cycle with standard chow and water provided *ad libitum.* Male and female mice between 12 and 24 weeks of age at the time of initial experimentation were used in the study. Only C57BL/6J male and female mice (Stock No. 000664, The Jackson Laboratory) were used.

### Female estrous determination

To determine the estrous stage of naturally cycling females used in sexual behavior experiments, vaginal smears were collected daily. Approximately 10 µL of sterile distilled water (dH₂O) was gently flushed into the vaginal canal several times using a 10 µL pipette tip, taking care not to insert the tip deeply to avoid inducing pseudopregnancy. The recovered cells were transferred onto a clean glass slide, air-dried, and stained with Mayer’s Hematoxylin (Sigma-Aldrich, Cat. MH516) for 1 min, followed by 0.1% Eosin Y (Sigma-Aldrich, Cat. 230251) in 95% ethanol for 30 s. Slides were rinsed with water, air-dried, and stored for later examination.

Vaginal cytology was assessed under bright-field illumination (10× objective) using a Leica light microscope. Estrous stages were identified based on predominant cell types: diestrus consisted of mostly leukocytes, proestrus was defined by mostly nucleated epithelial cells and some cornified epithelial cells, estrus was defined by cornified epithelial cells, and metestrus is defined by a mix of the three cell types (Byers et al. 2012).

In addition to cytological staging, visual inspection of the vaginal opening was performed as an independent validation (Byers et al. 2012). Briefly, each mouse was gently held by the tail with forepaws resting on the cage lid, and the vaginal opening was observed. Proestrus females exhibited a swollen, moist, and wide vaginal opening; estrus females had a less swollen but still open vaginal canal; metestrus and diestrus females showed closed or nearly closed canals.

### Ovariectomy (OVX) and hormonal priming of female mice

To standardize sexual receptivity in females paired with males (for male behavioral testing), OVX was performed followed by hormonal priming. The bilateral OVX procedure was adapted from Idris (2012). Female mice (≥ 6–8 weeks old) were deeply anesthetized with ketamine/xylazine (100/10 mg/kg, i.p.). After shaving and disinfecting the lumbar region, a single midline dorsal incision (∼ 0.5 cm in length) was made.

Subcutaneous connective tissue was separated from the underlying musculature, and a small incision was made on each side to enter the peritoneal cavity and expose the ovary. The ovarian fat pad and ovary were gently excised. In cases of bleeding, a single ligature was placed around the oviduct to minimize blood loss. The uterine horn and remaining oviduct were returned to the abdominal cavity, and muscle and skin layers were sutured. Mice received postoperative analgesia (meloxicam, s.c.). All procedures were performed under aseptic conditions.

Hormonal priming followed the protocol of Inoue et al. (2019). Estrus was induced in by delivering (subcutaneously) 10 µg of 17 β-estradiol benzoate (Sigma Aldrich, Cat. D9542 ) dissolved in 100 µL of sesame oil (Sigma Aldrich, Cat. S3547) two days prior to the mating experiment. 5 µg of 17 β-estradiol benzoate in 50 µL of sesame oil was delivered one day prior to the experiment, and 50 µg of progesterone (Sigma Aldrich, Cat. P0130) in 50 µg of sesame oil was given 3-6 hr before the mating experiment. Hormone-primed females were visually confirmed to be in estrus by observation of a wide vaginal opening.

### Locomotor activity (LMA) and body temperature (Tb) sensor surgery in male mice

For the serum testosterone measurements in males (described below), mice were implanted with LMA/Tb sensors. To achieve this, male mice (at least 6-8 weeks of age) were deeply anesthetized with a mixture of ketamine/xylazine (100/10 mg/kg, i.p.), as noted above. Radiotelemetry sensors (TA-F10, DSI, US) which provide both LMA and Tb data were implanted in the peritoneal cavity with a small abdominal incision. Mice receive analgesic treatment with meloxicam after surgery.

Average Tb and LMA were recorded every 5 min during all the protocols using the radiotelemetry DSI system. The signal from the telemetry probes were received and converted with the PhysioTel HD and PhysioTel (DSI) hardware. The LMA and Tb data were analyzed using the software ClockLab Analysis version 6 (Actimetrics). Tb and LMA were used to predict activity onset/offset in male mice housed in constant dark (DD) conditions.

### Male serum collection and testosterone measurements

Male mice (≥8 weeks of age) were implanted with Tb/LMA probes as described above. After recovery, each male underwent two separate mating experiences with OVX, hormone-primed female mice, with at least one week between sessions. After the second sexual experience, males were dark-adapted and allowed to free-run under constant darkness (DD) for at least two weeks prior to blood collection.

Samples (approximately 20–30 µL) were collected from tail vein blood at four circadian time points: CT1, CT7, CT13, and CT19. Samples were obtained every 30 hours to minimize stress and avoid anemia. After the fourth collection, mice were returned to a 12:12 LD cycle for at least two weeks before undergoing a second round of sampling at CT4, CT10, CT16, and CT22 (again, spaced by 30 hours between collections). Blood flow was stopped by applying gentle pressure with sterile gauze. All samples were collected within 30–60 seconds of initial handling to minimize handling-induced stress (Ramirez-Plascencia et al. 2025).

Blood was collected into microvette tubes (CB300; Sarstedt, USA) and centrifuged at 10,000 × g for 10 minutes at 4 °C. Serum was separated and stored at −40 °C until analysis. Testosterone concentrations were measured in duplicate using a commercially available ELISA kit (Crystal Chem, #80552), according to the manufacturer’s instructions. Samples with a coefficient of variation (CV) exceeding ±15 % between duplicates were excluded from analysis.

### Ultrasonic vocalization (USV) determination and analysis

Male USVs were captured at a 300 kHz sampling rate using an Avisoft UltraSoundGate 116H kit equipped with a condenser ultrasound microphone CM16/CMPA (Avisoft Bioacoustics Recorder; Berlin, Germany). This microphone was positioned approximately 5-6 inches above the mating arena.

Once captured by the Avisoft recording system, all USV files were transferred to Avisoft-SASLab Pro software (version 4.53) for spectrogram visualization and post-processing and analysis of USV number, duration, amplitude, and frequency. Detected calls were visually inspected to confirm accuracy prior to statistical analysis.

### Short-term mating paradigm

#### Male short-term mating experiments

Prior to short-term (20 min) mating experiments, male mice were provided with two sexual experiences to facilitate the development of mating behavior and improve sensory cue processing (Swaney et al. 2012) (Remedios et al. 2017). Males were group-housed until their first sexual experience, after which they were single-housed for the remainder of the study.

For the first sexual experience, each male was paired overnight with an OVX, hormone-primed, sexually experienced female. Hormone-primed females were given a prior mating round to ensure robust sexual receptivity, as females can exhibit reduced receptivity during their first mating encounter (Thompson and Edwards 1971; Xu et al. 2012). The following morning, females were removed and examined for the presence of a copulatory plug. After a minimum one-week interval, a second sexual experience was performed using a novel female. This second experience occurred at a randomly assigned time between ZT10-ZT14 in the experimental mating arena for 20 minutes, to habituate the male to experimental conditions (e.g. the buzz tone from the Arduino buzzer that synchronized video and audio, e.g. USV, recordings). It was not a requirement for the males to copulate with the female partners during the two sexual experience trials.

At least seven days were allowed between the last sexual experience and the first experimental session, given that male mice have a refractory period require approximately one week to recover from sexual satiety (Zhang et al. 2021). Two days before the experiment, males were dark-adapted to remove light-based timing cues.

During the experiment, males were randomly tested at one of four circadian timepoints (CT1, CT7, CT13, or CT19). A separate cohort was tested at CT4, CT10, CT16, or CT22 to avoid overexposure to sexual experience within a month-long period. Experiments were spaced seven days apart to control for the male refractory period.

For each mating session, the male remained in its home cage, which was placed inside a light-tight cabinet equipped with a USV microphone and camera. The environment was illuminated only by infrared light. Food and nesting materials were removed to avoid occlusion. Males were given 5 minutes to habituate, after which a novel OVX, hormone-primed female was gently introduced. Sessions lasted 20 minutes and were recorded with a Blackfly FLIR camera (30 Hz). USVs were synchronized with video recordings using an Arduino buzzer (tone ∼5 kHz), to which both sexes were habituated during their second sexual experience. Females were always mated at their zeitgeber time (ZT)13 to control for female behavioral variability.

A 5V transistor–transistor logic (TTL) pulse triggered the camera. After 20 minutes, the female was removed, checked for a copulatory plug, and the male was returned to its 12:12 LD cycle. The same procedure was repeated weekly at a different CT for three additional sessions, yielding four total circadian timepoints per male. Males that did not display mounting behavior at any timepoint were excluded from analysis.

### Female short-term mating experiments

Prior to testing, naturally cycling females underwent two sexual experiences during estrus, spaced at least one cycle apart. Each experience involved pairing an estrus female with a novel, sexually experienced male for up to 2 hours or until intromission was observed. During the second experience, both sexes were placed in the mating arena to habituate to the experimental setting. The experimenter observed each session, and mating was terminated immediately at intromission to prevent pregnancy. To rule out pseudopregnancy, at least 8–10 days elapsed between the final sexual experience and the experimental session. Vaginal cytology was monitored daily, as pseudopregnancy is associated with a prolonged diestrus phase. A total of 7 female mice did not cycle consistently and were not used further in the study. Females were dark-adapted to DD for two days in single housing before the experimental session.

During experiments, each female (in confirmed estrus based on vaginal smears taken 2-3 hours before the experiment or based on visual inspection) was mated only once, at one of eight circadian timepoints (CT1, 4, 7, 10, 13, 16, 19, or 22). Mating sessions were 40 minutes in duration in darkness, as preliminary trials indicated females required longer to display receptive behaviors. Females were paired with novel, sexually experienced males, each maintained on a LD schedule such that they were tested at their ZT16 (the male sexual behavior peak; Fig. 1). Each session took place in the same conditions described above for males. After 40 minutes, females were checked for a copulatory plug and returned to their LD cycle. No further mating was performed unless the paired male failed to mount, in which case the female was re-tested several cycles later with a different male.

### Long-term mating paradigm

For the long-term mating paradigm, we adapted the protocol from Kukino et al. (2022). 13 pairs of sexually experienced male and female mice were used. Females were naturally cycling and had prior sexual experience but had never been pregnant (previous mating sessions were terminated at intromission). Both sexes were maintained under a standard 12:12 h light–dark (LD) cycle and had not engaged in sexual activity for at least one week prior to the start of the experiment. To monitor estrous stage, vaginal smears were collected daily for five consecutive days leading up to the start of mating (including the day of pairing). Prior to the experiment, males and females were each habituated for 1 hour to the experimental cage and light-tight recording cabinet. The mating cage consisted of a clear 7-quart storage bin tall enough to prevent easy escape and was left uncovered during the experiment. Continuous video recordings were obtained using an XVIM 8- channel 1080P infrared (IR) camera system positioned above the cage.

Each pair was housed together continuously for six days, a duration chosen to capture at least one complete estrous cycle (average length 4–5 days) and to allow sufficient opportunity for sexual behavior and potential pregnancy. Mating sessions began at either CT1, CT6, or CT12 to assess whether the timing of pairing influenced sexual behavior dynamics. Because males often display heightened sexual activity upon initial pairing with a novel female, these different CT start times allowed assessment of potential circadian effects on mating onset. The entire 6-day long experiment was conducted in DD conditions, with illumination provided only by the IR lights from the recording cameras. No experimenter interventions occurred during the 6-day period. In two pairings, brief technical issues resulted in recording interruptions (1–2 hours). After six days, pairs were separated and returned to their original cages under standard LD conditions. Recorded videos were later manually analyzed for sexual behaviors (see Analyses section below), and females were subsequently monitored for parturition.

### Analyses of male and female mating behaviors

Videos from the short mating experiments were manually scored by an observer blind to experimental conditions. A *mount* or *mount attempt* was defined as the moment when the male climbed onto the female and began rapid, shallow thrusting, and it was considered terminated once the male dismounted. *Intromission* was identified when thrusting became slower and deeper, and *ejaculation* was noted when an intromitted male fell to the side and remained motionless for several seconds before grooming himself (Zhang et al. 2021). At the end of each session, females were checked for the presence of a copulatory plug to confirm successful fluid transfer.

Female sexual behaviors were also analyzed from these videos. *Receptive behavior* was defined as the female assuming a lordotic posture or not resisting male mount attempts. *Rejection behavior* was defined as any active resistance to mounting, including escaping during or prior to a mount, rising onto its hind legs, or kicking at the male. Because female sexual behavior depends on the number of male mounting attempts (Bonthuis et al. 2011), females whose partners exhibited fewer than five total mounts during the 40-minute session were excluded from analysis. The latency for the male to mount was subtracted from the latency for the female to exhibit rejection behaviors. The rejection and lordosis quotients were also calculated post-hoc. The rejection quotient was defined as the number of female rejections/total male mount attempts (x100). Similarly, the lordosis quotient was defined as the number of lordotic responses/total male mount attempts (x100).

Videos from the long-term (6-day) mating experiments were similarly analyzed by an observer blind to group assignment. Mating behaviors were classified as *unsuccessful mounts*, *successful mounts*, *intromission*, or *ejaculation*. *Unsuccessful mounts* were defined as attempts in which the female resisted (e.g., darted away, reared up, or kicked) during a male’s approach. *Successful mounts* were defined as instances when the female adopted a stable, lordotic posture or did not actively resist. To avoid double-counting closely spaced events, at least five seconds had to elapse between successive mounting attempts for them to be scored as separate bouts.

### Misalignment and alignment mating paradigm

Female mice were maintained on a 12:12 LD cycle, and vaginal cytology was performed daily to determine estrous stage, as described above. Male mice received two prior sexual experiences, spaced at least seven days apart, in which they were permitted to mate overnight with a novel, OVX, hormone-primed female. Similarly, female mice received two prior sexual experiences with a novel, sexually experienced male, during which mating was permitted up to the point of intromission or for two hours on the day of estrus.

Following these sexual experiences, male and female mice were individually housed and assigned to one of four experimental groups:

- Group 1 (Aligned Peak–Peak): Males mated at their circadian activity peak (e.g., CT16; see Fig. 1) with females also at their peak (e.g., CT16; see Fig. 2)
- Group 2 (Male Peak–Female Nadir): Males mated at CT16 with females at their CT4 nadir (see Fig. 2)
- Group 3 (Male Nadir–Female Peak): Males mated at their CT7 nadir (see Fig. 1) with females at CT16
- Group 4 (Aligned Nadir–Nadir): Both males and females mated at their circadian nadir (e.g., CT or CT4)

To achieve this circadian alignment or misalignment, all females were maintained on a standard 12:12 LD cycle and mated at either CT4 or CT16. Male mice were also housed on a 12:12 LD cycle, but the timing of lights-on and lights-off was shifted differently across the four male groups so that mating times were either aligned or misaligned relative to the female’s circadian phase. All mice were dark-adapted for two days prior to the mating experiment.

Vaginal cytology was used to track individual estrous cycles and predict the day of estrus. Females were only paired for mating when in confirmed estrus. Preliminary pilot experiments using a 40-minute mating session (similar to the short mating paradigm described above and in Fig. 2) resulted in successful copulation in only ∼20% of females. Therefore, for the main experiment, the mating period was extended to one hour. Females were allowed a second mating opportunity (several cycles later) with a novel male if intromission and/or ejaculation did not occur during the first session, reducing the total number of animals required for the study. If copulation did not occur during this second session, the female was not used for further mating sessions.

During each mating session, the male was gently introduced into the female’s home cage, and behavior was recorded using a Blackfly FLIR camera. Videos were live-streamed to a laptop to allow real-time observation of mating progress and ejaculation. After one hour, the male was removed and returned to his home cage.

Copulatory plugs were checked to validate ejaculation. Successfully inseminated females were returned to a standard 12:12 LD cycle and left undisturbed, except for ultrasound imaging at 6–7 days post-conception (described below). Pregnancy outcomes were monitored, and females were spot-checked every 3–6 hours during the predicted parturition window to document litter size, pup sex ratios, and birth weights.

### Ultrasound imaging

Following the 1-hour alignment and misalignment mating paradigm described above, females that had copulatory plugs and were observed to have successfully mated were returned to their home cages. Ultrasound imaging was performed 6–7 days post-coitum to determine pregnancy status and quantify gestational sacs, following protocols adapted from (Pallares and Gonzalez-Bulnes 2009). Ultrasounds were conducted between ZT8-ZT12 whenever possible. 6-7 days post coitum was chosen as the ideal time to image the female mice given that 5.5 days post-mating was the earliest time in which embryo vesicles and embryos could be detected (Pallares and Gonzalez-Bulnes 2009), and we wanted to determine gestational sac number prior to complete resorption occurring (Drews et al. 2020).

For imaging, females were anesthetized with 3–4% isoflurane in 1–1.5 L/min oxygen for induction and maintained at 1–2% isoflurane via a nose cone during the procedure. Respiratory and heart rates were continuously monitored by visual inspection throughout the session. Each mouse was positioned supine on a heated platform and secured gently with adhesive tape. Abdominal fur was removed using Nair Body Cream, and a conductive gel (Spectra 360 electrode gel; Parker Labs, Product No. 12-08) was applied to eliminate air gaps between the transducer and the skin surface.

Ultrasound examinations of gestational sacs were conducted using a VisualSonics Vevo 2100 system equipped with an MS500S transducer (40 MHz). The transducer was gently moved across the abdomen to visualize the uterine horns, using the bladder and cervix as anatomical landmarks. The presence and number of gestational sacs were recorded for each uterine horn, and sac lengths were measured when possible. Each imaging session lasted approximately 5 minutes per animal.

No adverse effects were observed in females undergoing ultrasound imaging. Following the procedure, mice were placed on a heating pad for at least 10 minutes before being returned to their home cages. The transducer and imaging platform were cleaned between animals to prevent cross-contamination.

### Validation of ultrasound imaging

The accuracy for detecting very early pregnancy (and gestational sac number) was validated in four pregnant female mice that, after ultrasound imaging (at 6 days post-coitus), were immediately euthanized (via CO₂ inhalation). The uterine horns and ovaries were removed and the number of gestational sacs/embryos were counted. Additionally, 3 non-mated (and therefore non-pregnant) females were also imaged via ultrasound so that the experimenters could accurately determine the presence vs absence of gestational sacs. 2 non-pregnant females were euthanized, and uterine horns were removed, inspected, and as expected, no gestational sacs/embryos were present.

For all ultrasounds (images and videos) were evaluated by the initial ultrasound operator, and if they did not show obvious gestational sacs at the 6-7 day ultrasound, were also validated by a second blinded observer for completeness.

### Gestational sac measurements

The diameter of gestational sacs at 6-7 days post coitum were measured using the Fujifilm VisualSonics Vevo LAB software suite, which includes a package for embryonic measurements. A linear measurement tool was used to determine the diameter of each gestational sac, and a blind experimenter took the measurements directly on the ultrasound image. Importantly, only those gestational sacs that were clearly visible in the ultrasound videos and images were analyzed. Thus, embryos that were observed (live) during ultrasound imaging, may not have been measured post-hoc if the image was not clear. Similar to Szalai et al. (2015), gestational sac size ranged between ∼0.9 mm and ∼2.8 mm. This range likely reflects biological variability in sac size, timing of mating, and whether the ultrasound was conducted at 6 vs 7 days post coitum.

### Statistical analysis

Statistics were performed using GraphPad Prism 10 software, and all data is presented as the mean ± SEM. Statistical significance for all experiments was set at p < 0.05 (*). Power analysis for the short mating paradigm experiments was conducted using preliminary data and prior experimental results from the lab to detect an effect size of 0.8 (Cohen’s d) with α = 0.05 and power = 0.8. Statistical comparisons between two groups were performed using Student’s t-tests, while comparisons between more than 3 circadian timepoints (or groups) were done using an ANOVA. Post-hoc tests were conducted to show an interaction obtained from significant ANOVA results. To test whether early embryonic size could predict pregnancy outcome, logistic regression was performed relating average gestational sac diameter to term pregnancy success. Odds rations, confidence intervals, and model fit were evaluated as part of this analysis. Finally, Grubb’s tests was performed on each data set to test for statistically significant data points/outliers in each group. For analysis of the long-term mating experiment, for each animal, graphs were created by binning event times into 24-hour bins and normalizing the bins to yield a probability distribution (sum = 1). Normalized distributions were then averaged across animals to obtain the population-level temporal probability profile. Representations were then created in Matplotlib Python Library or MATLAB.

## Acknowledgements

This study was supported by NIH Grants: 1F32HD111339-01A1 to SA, T32HL007901 to SA, 2023 Focused Projects Grant for Junior Investigators; American Academy of Sleep Medicine to SA, and R01NS072337 to CBS.

The authors thank Quan Ha and Sathyajit Bandaru for their superb technical assistance. The authors also thank Dr. Carrie Mahoney for the thoughtful conversations and suggestions, and Dr. Stephen Zhang for assisting with the mating experimental setup. They also thank Dr. Sean Clohessey (and lab) for assisting with the ultrasound setup.

---

**Supplementary Figure 1:**
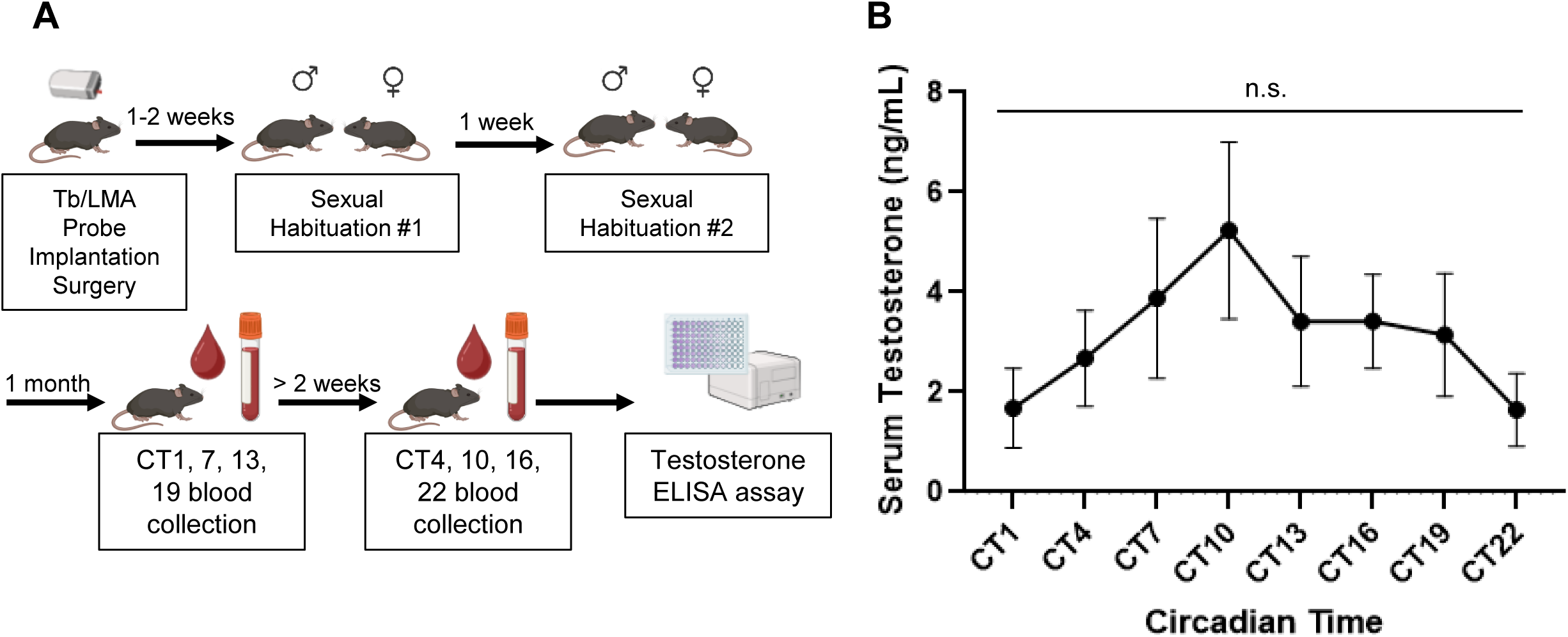
Assessing testosterone levels across the circadian day. A) Male mice were implanted with body temperature/locomotor activity abdominal probes. After a 1-2 week recovery from surgery, they underwent their first overnight sexual habituation with an OVX, hormone-primed female. The second overnight sexual habituation occurred about 1 week later. At least 2 weeks after the second sexual habituation, male mice were dark adapted for at least 2 weeks. After being in DD for at least 2 weeks, blood collections began. Blood was collected from the tail vein every 30 hr beginning at CT1 and continuing at CT7, CT13, and CT19. After the CT19 blood draw, mice were returned to LD for at least 2 weeks before being returned to DD for CT4, CT10, CT16, and CT22 blood sampling to begin. Plasma from the blood collections was then used for a testosterone ELISA assay. B) Testosterone peaked at CT10, although significance was not reached. n.s.: not significant. N = 7-10 male mice per timepoint.

